# Systematic detection of Mendelian and non-Mendelian variants associated with retinitis pigmentosa by genome-wide association study

**DOI:** 10.1101/859744

**Authors:** Koji M Nishiguchi, Fuyuki Miya, Yuka Mori, Kosuke Fujita, Masato Akiyama, Takashi Kamatani, Yoshito Koyanagi, Sato Kota, Toru Takigawa, Shinji Ueno, Misato Tsugita, Hiroshi Kunikata, Katarina Cisarova, Jo Nishino, Akira Murakami, Toshiaki Abe, Yukihide Momozawa, Hiroko Terasaki, Yuko Wada, Koh-Hei Sonoda, Carlo Rivolta, Tatsuro Ishibashi, Tatsuhiko Tsunoda, Motokazu Tsujikawa, Yasuhiro Ikeda, Toru Nakazawa

## Abstract

To uncover genetic basis of autosomal recessive retinitis pigmentosa (ARRP), we applied 2-step genome-wide association study (GWAS) in 640 Japanese patients prescreened with targeted re-sequencing. Meta-GWAS identified three independent peaks at *P* < 5.0×10^-8^, all within the major ARRP gene *EYS*. Two were each tagged by a low frequency variant (allele frequency < 0.05); a known founder Mendelian mutation (c.4957dupA, p.S1653Kfs*2) and a presumably hypomorphic non-synonymous variant (c.2528G>A, p.G843E). c.2528G>A newly solved 7.0% of Japanese ARRP cases, improving genetic diagnosis by 26.8% and simultaneously serving as a new attractive target for genome editing gene therapy. The third peak was tagged by an intronic common variant, representing a novel disease-susceptibility signal. GWAS successfully unraveled genetic causes of a rare “monogenic” disorder for the first time, which provided unexpected insights into significant contribution of non-Mendelian genetic factors and identified a novel high frequency variant directly linked to development of local genome therapeutics.

## Introduction

Genetic diagnosis of heterogenous inherited disorders became less challenging after the wide availability of next generation sequencing. However, although the technological development has substantially improved diagnosis rates, the genetic basis of disease remains unknown in a large proportion of patients, highlighting the limits of the simplex sequence-based approach. Retinitis pigmentosa (RP), which lacks effective treatment options, is the most common form of inherited retinal degeneration. It is initially characterized by the loss of rod photoreceptors, which mediate night vision, and then involves the loss of cone photoreceptors, which are responsible for daylight vision. RP affects approximately 1 in 3,000 people worldwide. The disease is highly heterogenous presenting with a various hereditary pattern^1^ ranging from classical Mendelian inheritance to non-Mendelian inheritance due to incomplete penetrance^2, 3^, hypomorphic allele^4, 5, 6, 7, 8^, or oligogenecity^4, 5, 9, 10^. However, despite numbers of reports of non-Mendelian inheritance causing RP, its significance in the context of overall genetic pathology of the disease is yet to be demonstrated. In Japan, the genetic basis of RP remains unknown in up to 70% of cases even after targeted re-sequencing, whole-exome sequencing, or whole-genome sequencing ^4, 11, 12, 13^. There is an urgent need for genetic diagnosis of these unsolved cases, particularly those with the autosomal recessive (AR) inheritance pattern of RP (ARRP), caused by loss-of-function mutations, because such patients may be amenable to adeno-associated virus (AAV)-mediated gene supplementation therapy^14^. Furthermore, there is a growing interest in the detection of prevalent founder mutations, which are also potential targets of AAV-mediated genome-editing therapy^15, 16, 17, 18^. These cases are also candidates for antisense oligo therapy^7, 16, 19^, which allows local treatment of the retinal genome and can target larger genes that cannot be treated with conventional gene supplementation therapy.

Genome-wide association study (GWAS) is a type of analysis that is most often applied to identify susceptibility loci for common traits, each with a relatively small genetic influence^20^. However, it can also uncover rare variants with strong genetic effects in complex diseases that behave almost as Mendelian alleles in “monogenic” diseases^21, 22, 23^. By comparing differences in allele frequency in cases and controls, GWAS can provide unbiased means of detecting disease-associated loci evenly across the genome preferentially in the order of disease contribution, with little assumption of the inheritance pattern. This contrasts with case-oriented sequencing approaches, which are often obliged to focus around exons and their boundaries, to identify mutations that follow classic Mendelian inheritance. Thus, GWAS can in theory complement these widely used sequencing approaches by searching for any significant genetic risks that remain undetected. However, GWAS has never been used to directly search for genetic risks in rare “monogenic” diseases, and its usefulness in such purposes remains unknown.

Here, we report the detection of three disease-associated signals/variants in patients with presumed ARRP, using an approach that combined GWAS with targeted re-sequencing.

## Results

### Detection of the *EYS* locus with GWAS and targeted re-sequencing

We gathered a total of 944 DNA samples from unrelated patients who had RP consistent with the AR mode of inheritance, including isolated cases with no family history, and 920 control samples. All samples were genotyped with a single nucleotide polymorphism (SNP) array. To search for undetected genetic risks contributing to ARRP, we carried out a meta-GWAS using two independent data sets (Table S1). Of the 644 cases and 620 controls genotyped in the first GWAS, 432 cases and 603 controls were used for analysis after removing 63 cases and 21 controls that failed quality control (QC) and an additional 149 “solved” cases in which targeted re-sequencing identified pathogenic mutations to account for the cause of disease^11^. Similarly, after removing 14 cases and 13 controls that failed QC and excluding 78 cases genetically solved with targeted re-sequencing^11^, the second GWAS included 208 cases and 287 controls. The results of the two GWASs are summarized in Figure S1 and Tables S2 and S3. Then these two GWAS datasets were combined (for a total of 640 cases and 890 controls) to carry out a meta-GWAS (Figure 1A). In this analysis, only the locus encompassing *EYS*, the most frequent ARRP-associated gene in Japanese patients, remained significant (OR = 3.95, *P* = 1.18×10^-13^). We observed 12 other peaks with *P* < 1.0×10^-5^ in which no known RP genes were included (Table S4). Subsequent conditional analysis of the *EYS* locus detected 3 independent genome-wide significant signals (*P* < 5.0×10^-8^; Peaks 1 - 3 in the order of significance, Table 1 and Figure 1B). We then checked the associations of the most significant variants (lead variants), tagging each locus and non-synonymous and splice site variants in the 640 cases using the targeted re-sequencing results^11^. While Peak 1 and Peak 3 were each linked to a low frequency variant (AF < 0.05), Peak 2 was associated with a common nonsynonymous variant. More specifically, Peak 1 (rs76960013, allele frequency (AF) = 0.0414, odds ratio (OR) = 3.95, *P* = 1.18×10^-13^) was in linkage disequilibrium (LD; *r*^2^ = 0.68) with c.2528G>A (p.G843E; hereafter termed G843E; Table 1). G843E with an AF (0.0171) unusually high for ARRP has been described in conflicting ways in past reports, as having uncertain significance ^23^, being non-pathogenic ^24^, possibly being pathogenic (although without sufficient supporting evidence)^25^, and was unreported in the two largest genetic screening projects targeting Japanese RP patients^9, 11^. The second peak, with much higher allele frequency and lower OR (Peak 2; rs59178556, AF = 0.2161, OR = 1.83, P = 3.79 ×10^-10^), was in strong LD (*r*^2^ = 0.97) with a common nonsynonymous variant, i.e., c.7666A>T (p.S2556C; hereafter termed S2556C variant; Table 1) registered as benign/likely benign in the ClinVar (https://www.ncbi.nlm.nih.gov/clinvar/). Peak 3 (rs79476654, AF = 0.0005, OR = 16.46, *P* = 2.45×10^-8^) was in LD (*r*^2^ = 0.78) with c.4957dupA (p.S1653Kfs*2) (hereafter termed S1653Kfs; Table 1), recognized as a founder AR mutation^11^. It was detected even after removing solved cases with biallelic *EYS* mutations (including homozygotes and compound heterozygotes with S1653Kfs) screened by target re-sequencing prior to GWAS, because a large number of heterozygous carriers of S1653Kfs mutation remained genetically unsolved^9, 11, 24^. Haplotype analysis of *EYS* based on SNP array data and the results of targeted re-sequencing in RP patients confirmed that none of the lead variants of the identified signals were in LD with c.C8805A (p.Y2935X) or c.G6557A (p.G2186E) (hereafter termed G2186E), the two other known founder mutations in this gene^9^, in contrast to S1653Kfs, which was in LD with Peak 3 (Figure 1C and Tables 1 and S5). This suggests that Peaks 1 and 2 represent under-recognized genetic risks in *EYS*. Thus, GWAS in combination with targeted re-sequencing successfully detected disease-associated variants overlooked by simple sequence-based approaches in a rare “monogenic” disease.

**Figure 1.**
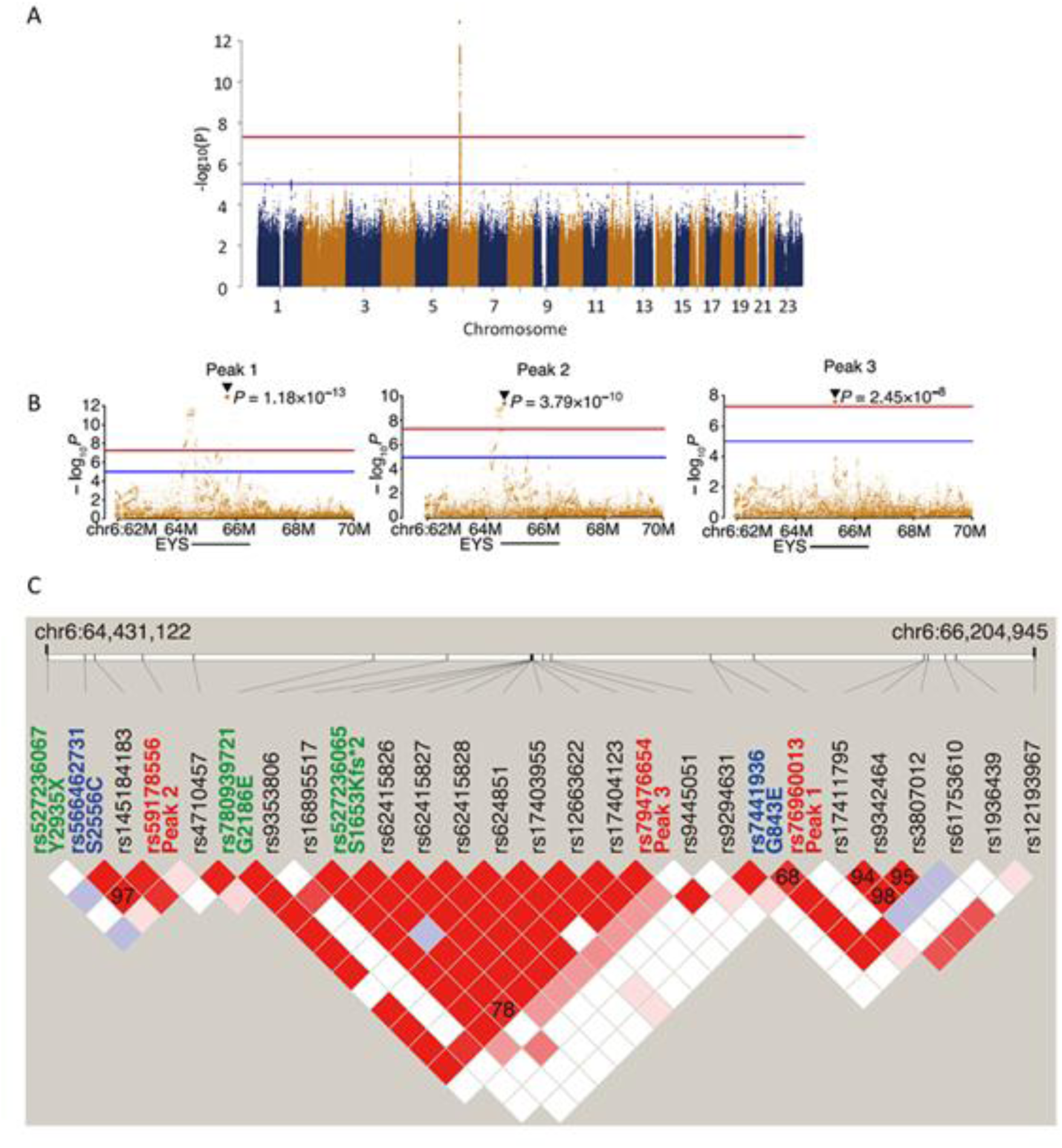
Genome wide association study (GWAS) of presumed ARRP patients. **A.** Results of a meta-GWAS displayed as a Manhattan plot. Genome-wide significance (*P* = 5.0 ×10^-8^) and possible significance (*P* = 1.0×10^-5^) are marked with red and blue lines, respectively. A single peak at the *EYS* locus surpassed genome-wide significance. **B.** Results of a conditional analysis presented as a regional plot. Three independent peaks at P < 5.0×10^-8^ were delineated after conditioning (Peaks 1-3). **C.** LD plot using all non-synonymous variants (identified in > 5% of cases) and lead variants for Peaks 1-3 identified in GWAS in presumed ARRP patients. The LD plot was generated using Haploview (ver. 4.1). The default color setting of the software was used for block color setting (D’ / LOD). The numbers on the blocks indicate *r^2^* x 100; numbers are shown on the blocks only for pairs with *r^2^* >0.3. Peaks 1, 2, and 3 were in LD with G843E, S2556C, and S1635Kfs, respectively. The lead variants for Peaks 1-3 are shown in red. Reported pathogenic founder mutations^11^ are shown in green, while non-synonymous variants linked to the lead variants are shown in blue.

**Table 1.** Peaks detected with conditional analysis of the *EYS* locus and associated nonsynonymous variants. Information on the 3 independent peaks detected in this study and exonic variants in LD (shaded) are presented. *The odds ratio for S1653Kfs was not available (NA), because the variant was not included in the imputed genotypes of the GWAS analysis.

|  | Peak 1 | Peak 2 | Peak 3 |
| --- | --- | --- | --- |
| dbSNP rsID | rs76960013 | rs59178556 | rs79476654 |
| Gene info | EYS:intron | EYS:intron | EYS:intron |
| Ref/Alt | A/G | C/A | T/C |
| ToMMo AF | 0.0414 | 0.2161 | 0.0005 |
| Odds ratio | 3.95 | 1.83 | 16.46 |
| *P* value | 1.18E-13 | 3.79E-10 | 2.45E-08 |
| Linked variant | G843E | S2556C | S1653Kfs |
| dbSNP rsID | rs74419361 | rs66462731 | rs527236065 |
| ToMMo AF | 0.0171 | 0.214 | 0.0044 |
| Odds ratio | 3.75 | 1.79 | NA* |
| Corrleation ($r^2$) | 0.68 | 0.97 | 0.78 |

### Expression analysis of G843E allele in genome-edited patient-derived lymphoblasts

The allele frequency (0.2140) of S2556C, linked to Peak 2, was undoubtedly too high for a pathogenic Mendelian mutation causing a rare “monogenic” disease. On the other hand, the allele frequency of G843E linked to Peak 1 was much lower (0.017), yet still too high for a classical AR allele, raising the possibility that it represents a nonpenetrant or hypomorphic ARRP allele. However, it is also possible that a true ARRP mutation in LD with Peak 1 exists deep in the non-coding region. However, the vast majority of the pathogenic mutations in *EYS* are either nonsense, frameshift, or slice site mutations^11^ that would presumably result in a qualitative alteration in the mRNA sequence. Thus, we directed our search to mainly variants that could affect splicing. For this purpose, we carried out two experiments. First, we performed whole genome sequencing (WGS) in two G843E homozygotes and two compound heterozygotes (G843E and S1653Kfs or G2186E). We found that there were no obvious structural variants in *EYS* that affected coding sequence. A splice site prediction analysis^24^ also detected no coding and non-coding variants that could alter splicing in these patients. Second, we established patient-derived lymphoblasts from homozygotes of G843E and S1653Kfs and studied the expression of *EYS* mRNA by forced transcription of *EYS* through insertion of a constitutively active *CAG* promoter immediately upstream of the initiation codon of the gene (Figure S3)^15^. Among the seven main transcript variants reported^25^, the retina-specific long isoforms (transcription variants 1 and 4) are considered essential for photoreceptor biology^25, 26^. RT-PCR followed by Sanger sequencing indicated that, mRNA containing G843E was expressed without the loss of the C-terminal end of the retina-specific long isoform (Figure 2A and B). This was unlike transcripts with homozygous S1653Kfs that resulted in the loss of this isoform via nonsense-mediated decay, which was successfully rescued by replacing the mutation by wildtype sequence through genome editing (Figure 2C). These results favor against the presence of an intronic mutation in LD with Peak 1 that results in altered splicing and a presumed premature termination of the reading frame, but support G843E as the causal mutation linked to Peak 1.

**Figure 2.**
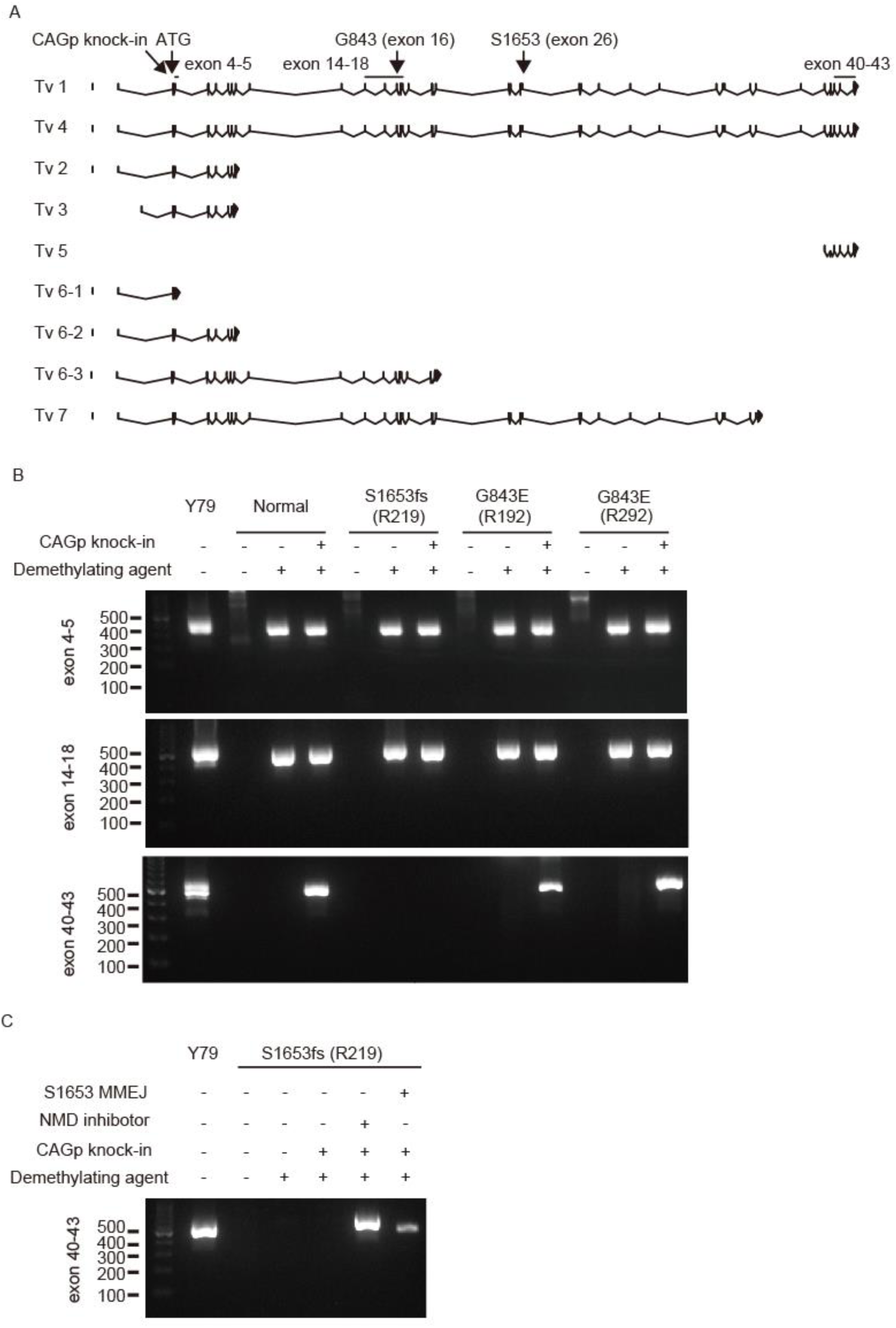
Expression analysis of the *EYS* G843E mutant allele in patient-derived lymphoblastoid cell lines. **A**. A schematic map of the RT-PCR primer designed in relation to the exon-intron structure and mutations (G843E and S1653Kfs) in *EYS* and published transcript variants (Tv).^25^ The locations of G843E (Exon 16) and S1653Kfs (Exon 26) are indicated by the arrows. Exon numbers are based on Tv1. Note, Tv5 was identified only in fibroblasts^25^. **B**. RT-PCR analysis of *EYS* expression. The regions for exons 5–6, exons 14–18, and exons 40–43 of *EYS* were amplified on cDNA generated from patient-derived lymphoblast cell lines with wildtype *EYS* (normal), homozygous S1653Kfs, and homozygous G843E. The Y79 retinoblastoma cell line was used as a positive control. Note C-terminal exons the long isoform Tv1 were detected in LCLs with homozygous G843E but not in that with homozygous S1653Kfs. Sanger sequencing of the RT-PCR amplicon confirmed the expression of the G843E mutation using a primer pair targeting exons 14-18. **C**. RT-PCR analysis of the lymphoblast cell line after mutation replacement genome editing treatment (MMEJ) or inhibition of nonsense-mediated mRNA decay (NMD) in S1653Kfs homozygote, after which expression of exons 40-43 was detected. MMEJ, microhomology-mediated end joining.

### Functional analysis of *EYS* G843E in zebrafish

Among mammals, only primates have *EYS* gene. And zebrafish (Danio rerio) is the only model in which loss-of-function mutations in the homologous eys has been shown to recapitulate photoreceptor degeneration observed in RP patients with *EYS* mutations^27, 28, 29^. As expression of G843E mutant in the mRNA has been confirmed with the patient-derived LCL (Figure 2), we used zebrafish to directly assess the function of the mutant allele. Endogenous Eys protein localized near the basal interface of the connecting cilium of the photoreceptors in adult fishes (Figure 3A, B). During development, Eys expression was observed after 4 days post-fertiliztion (dpf, Figure 3D-F). Morpholino-mediated knockdown of *eys* resulted in prominent rhodopsin mislocalization, a hallmark of photoreceptor degeneration^30^, at 7 dpf (Figure 3G,H). This abnormality was reversed more evidently by injection of wildtype human *EYS* mRNA (transcript variant 1; Figure 3J) compared to the mutant counterpart with G843E variant (Figure 3I, K). These results provide direct evidence for dysfunction of *EYS* caused by G843E.

**Figure 3.**
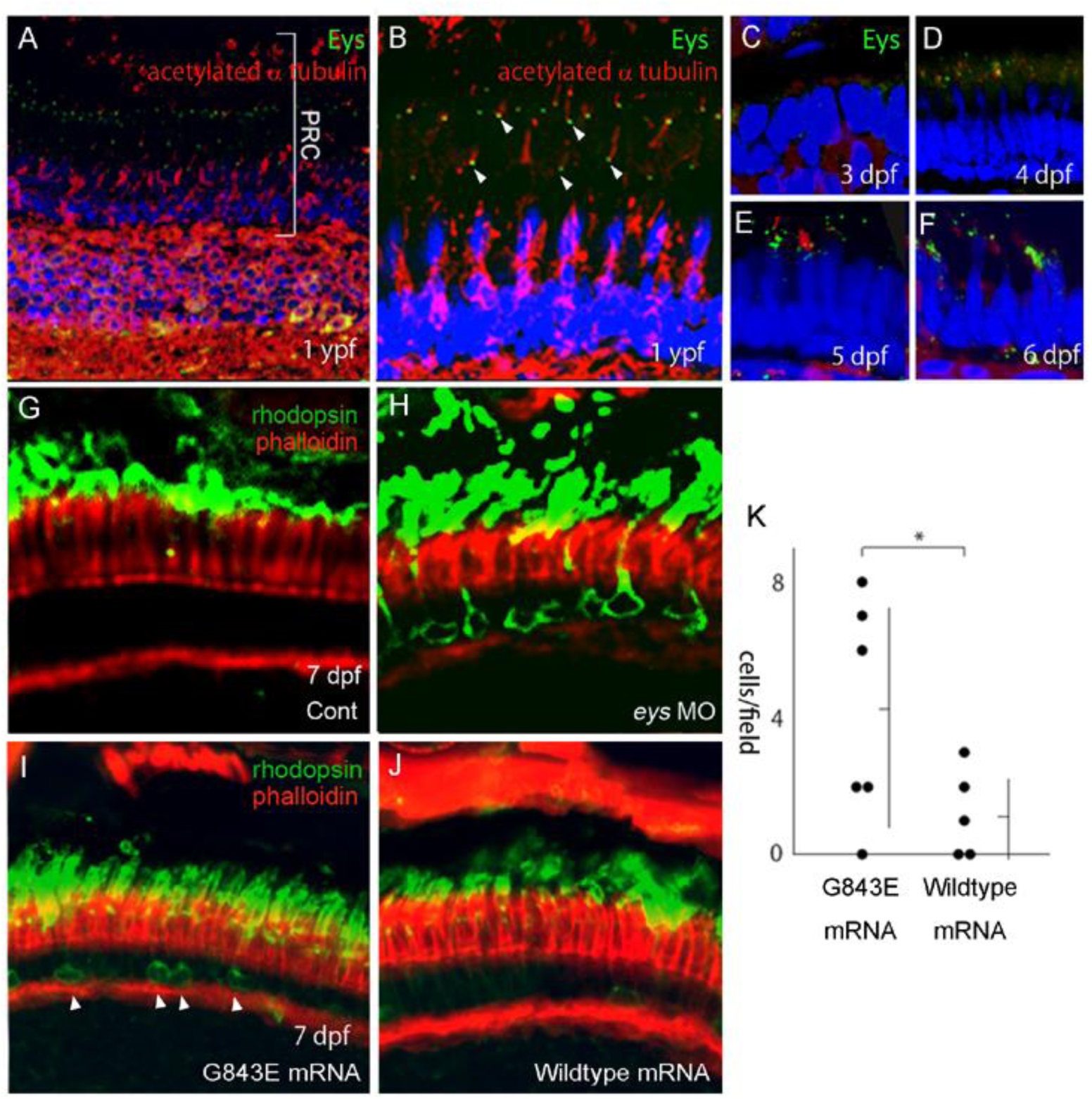
Functional analysis of *EYS* G843E variant in zebrafish. **A.** Immunostaining of Eys (green) in zebrafish retina at 1 year post-fertilization (ypf). **B**. High magnification image of photoreceptors. Eys (arrowhead) localized at the basal side of connecting cilium (acetylated α tubulin, red) of the photoreceptors. **C-F**. Expression of Eys during development at 3 days post-post-fertilization (pdf), 4 dpf, 5 dpf and 6 dpf. **G-H**. Rhodopsin localization in the photoreceptors at 7 dpf. **G**. Rhodopsin is localized at the photoreceptor outer segments in the control. **H**. *eys* knockdown by morpholino (MO) induced rhodopsin mislocalization toward the basal and the lateral membrane of the photoreceptors. **I**,**J**. Greater improvement of the rhodopsin mislocalization was achieved in the eyes supplemented with wildtype human *EYS* mRNA (**I**) over those injected with mutant human *EYS* mRNA (G843E; **J**) after MO-mediated knockdown of *eys*, indicating decreased *EYS* function by the mutation. **K.** A quantitative analysis of **I** (N = 5) and **J** (N = 6). Numbers of cells with mislocalized rhodopsin in the field were counted (vertical bar: mean ± standard deviation). *p < 0.05 (Unpaired two-tailed *t*-test). PRC, photoreceptors.

### Enrichment of G843E in genetically unsolved heterozygous carriers of another *EYS* mutation

A recent large-scale mutation screening project in 1,204 Japanese RP cases revealed an unusually high frequency of carriers of heterozygous deleterious mutations in *EYS,* accounting for 25.1% of the unsolved cases^11^, strongly indicating that there are AR mutations in *EYS* yet to be identified. Keeping this in mind, when we specifically looked at RP patients who were still genetically unsolved after targeted re-sequencing^11^, we found that G843E was highly enriched in patients with a heterozygous deleterious mutation in *EYS* (allele frequency 17.0%) compared to those without (allele frequency 6.9%, odds ratio = 2.46, *P* = 8.51×10^-7^, Fisher exact test; Table 2) or to the general population using a public database (allele frequency 1.7%, odds ratio = 10.0, *P* = 2.21×10^-32^, Fisher exact test; Table 2). This establishes that the G843E allele contributes to RP in *trans* with another *EYS* mutation, as in ARRP. Similarly, the frequency of G843E homozygotes was significantly higher (odds ratio = 97.0, *P* = 9.89×10^-12^) in genetically unsolved RP patients (13/640) compared to the general population (1/4773) establishing that the G843E allele contributes to RP in homozygosity as well, typical for an ARRP mutation. Meanwhile, analysis of Peak 2, linked to S2556C, also revealed significant enrichment of the variant in unsolved patients with a heterozygous deleterious mutation in *EYS* compared to those without (*P* = 2.56 ×10^-7^, Fisher exact test), although the difference was relatively small (allele frequency 39.1% vs 31.2%, odds ratio = 1.25). Taken together, the G843E mutation may cause RP when both alleles of *EYS* are affected, either in a compound heterozygous or a homozygous state, as observed in an ARRP allele. Meanwhile, Peak 2 may to confer a distinct pathomechanism.

**Table 2.** Analysis of target re-sequencing data of G843E in *EYS*. ToMMo: Normal Japanese population (N = 4,773). EYS: RP patients genetically solved with biallelic *EYS* mutations. nonEYS: RP patients genetically solved with mutations in genes other than *EYS*. with EYS path mut: Genetically unsolved RP patients with a heterozygous deleterious *EYS* mutation. with EYS path mut: Genetically unsolved RP patients with no deleterious *EYS* mutation. *EYS* mutations were extracted from a previous targeted re-sequencing study. ^11^

|  |  | Controls | Solved cases (N=227) |  | Unsolved cases (N=640) |  |
| --- | --- | --- | --- | --- | --- | --- |
|  |  | ToMMo | EYS | nonEYS | with EYS path mut | no EYS path mut |
| Genotype count | REF/REF | 4,611<br>(96.6%) | 76<br>(96.2%) | 139<br>(94.6%) | 98<br>(66.7%) | 437<br>(88.6%) |
|  | REF/G843E | 161<br>(3.4%) | 3<br>(3.8%) | 8<br>(5.4%) | 48<br>(32.7%) | 44<br>(8.9%) |
|  | G843E/G843E | 1<br>(0.0%) | 0<br>(0.0%) | 0<br>(0.0%) | 1<br>(0.6%) | 12<br>(2.4%) |
| Allele count | REF | 9,383<br>(98.3%) | 155<br>(98.1%) | 286<br>(97.3%) | 244<br>(83.0%) | 918<br>(93.1%) |
|  | G843E | 163<br>(1.7%) | 3<br>(1.9%) | 8<br>(2.7%) | 50<br>(17.0%) | 68<br>(6.9%) |
| MAF |  | 0.017 | 0.019 | 0.027 | 0.170 | 0.069 |

### Segregation analysis

Although G843E is consistent with an ARRP variant according to the analysis above, it is unlikely that the G843E allele acts as a simple Mendelian allele, considering its relatively high allele frequency in the general population (1.7%). Theoretically, even homozygotes of G843E alone would cause ARRP in at least 1 in 3,460 births, with a modest assumption of random mating, which is more frequent than the reported overall prevalence of ARRP in Japan (1 in 7,000)^31^. Furthermore, although the allele frequency of G843E (1.7%) is 3.8-fold higher than that of the founder variant S1653Kfs (0.44%) in the general population (Table 1), the observed frequency of homozygotes of G843E (14/867) is actually lower compared to that of S1653Kfs (24/867). This could be accounted for by G843E causing incomplete penetrance or a mild retinal phenotype, both of which could lead to a large underestimation of disease frequency. To explore this possibility, we carried out a segregation analysis in 18 unaffected (and 1 affected) siblings of index patients with G843E (either in a compound heterozygous or a homozygous state, 13 families; Figure S4). None of the unaffected siblings of the patients carried biallelic *EYS* mutations, except for the brother of YWC133, who was unexpectedly found to be compound heterozygous for G843E and S1653Kfs. This 75-year-old man was considered unaffected according to a local ophthalmologist who had carried out cataract surgeries on both eyes within the preceding year. Re-assessment of the patient at Tohoku University Hospital revealed a mildly but clearly constricted visual field, accompanied by moderate attenuation of the retinal vessels and diffuse alteration of the retinal pigment epithelium with modest retinal thinning in both eyes, although he had normal visual acuity (20/20). Nevertheless, the marked reduction in the electrical response of the patient’s retina to light stimuli probed by electroretinogram indicated that he also had a mild form of RP. Thus, the results are consistent with G843E being a hypomorphic ARRP allele and show that it can indeed sometimes cause mild retinal disease that may be overlooked without a thorough assessment. This may partly account for the gap between the known prevalence of RP and the allele frequency of G843E, complicating its interpretation in the past^11, 13, 32, 33^. Assuming that G843E is an ARRP allele, the mutation would account for an additional 7.0% of Japanese cases of RP, which would increase the proportion of genetically solved cases by 26.8%, either as compound heterozygotes or homozygotes.

## Discussion

Although previous reports have used GWASs to identify rare penetrant pathogenic variants in complex diseases^21, 22, 23^, our study is the first to demonstrate that GWAS, with the help of targeted re-sequencing, can be applied effectively to identify genetic risks in heterogenous “monogenic” disorders. We successfully identified three independent disease-associated signals, all in the gene *EYS*, including a locus in LD with the known commonest founder mutation S1653Kfs in *EYS* that causes ARRP^11^. This confirmed the quality of GWAS and its ability to effectively detect classical Mendelian mutations. More important, we detected a locus in LD with G843E, a controversial variant that did not previously fulfill the standard criteria required to determine pathogenicity^11, 13, 33, 34^. Analysis of the targeted re-sequencing data revealed that G843E was highly enriched in heterozygous carriers of another deleterious *EYS* mutation and homozygotes compared to the general population, consistent with the allele mediating the AR mode of inheritance. Yet, the relatively high allele frequency of G843E contradicts the known prevalence of ARRP. A segregation analysis identified an elderly asymptomatic patient who was compound heterozygous for G843E and S1653Kfs and had been erroneously assigned as unaffected, probably based on a lack of symptoms or typical features of RP. This was consistent with G843E being hypomorphic sometimes causing very mild phenotype later in life. In such instances, the disease may be overlooked without an assessment by electroretinogram, the most sensitive measure to detect RP. Nevertheless, the strong evidence from the segregation analysis (P < 0.01), the presence of G843E *in trans* with an established pathogenic variant in multiple families, along with *in vitro* expression and *in vivo* functional analyses supporting dysfunction of G843E have allowed us to reclassify the variant as “pathogenic” according to the standard guidelines^34, 35^. G843E as a quasi-Mendelian variant will likely enable genetic diagnosis in an additional 7.0% of Japanese patients with ARRP, which would represent a 26.8% improvement in the diagnosis rate. At the same time, the importance of this finding extended far beyond the context of genetic diagnosis as a detection of a founder mutation with an extremely large disease contribution provides a unique opportunity for development of an AAV-mediated mutation replacement genome editing gene therapy, which has shown promising *in vivo* outcomes^15, 16, 36^. This demonstrates the robustness of the approach, considering that mutations in novel RP genes, which are continuously discovered by sequencing, rarely account for more than 1% of cases and are unlikely to be suitable targets for drug development.

Recently, enrichment of the G843E variant in *EYS* in a group of patients with hereditary retinal degenerations (HRD) that carried a quasi-Mendelian allele in another gene (c.5797C>T/p.R1933* in *RP1*), has been reported^4^, suggesting indeed a non-Mendelian, oligogenic role of *EYS* in retinal degeneration. In our study, *RP1*-R1933* was infrequent among carriers of *EYS-*G843E, but this maybe attributable to gross differences in the clinical phenotypes considered (macular degeneration or cone-rod dystrophy in the *RP1* report^4, 37^ vs. canonical ARRP, studied here) Furthermore, while in heterozygous carriers *RP1*-R1933* seems to exert its pathogenic functions via the co-presence of *EYS*-G843E and other hypomorphic alleles outside of the *RP1* locus^4^, a reciprocal mechanism is not forcibly true, since molecular pathology of *EYS*-G843E in ARRP may follow different routes, as clearly shown above. Taken together, these results emphasize the unexpected pleiotropic role of *EYS*-G843E with respect to the range of unconventional genetic influence and its effect on clinical phenotypes.

GWAS also identified a novel RP-associated *EYS* signal (Peak 2) with no rare exonic or splice site variants in LD that could account for ARRP. It is possible that another quasi-Mendelian mutation in LD with Peak 2 in the non-coding regions remains undetected after targeted re-sequencing^4, 7, 19^. However, the higher frequency of the lead variant (AF = 0.216) for this peak is distinct from those of the other peaks (AF = 0.041 and 0.0005), resulting in a lower OR (1.83) well within the range of those for more common retinal diseases^38^. Therefore, it is possible that the true pathogenic variant(s) in LD may be high frequency variant(s) behaving in a non-Mendelian manner, similar to those presumed to account for susceptibility signals in common diseases. Although, the exact mode of genetic influence remains to be elucidated for this peak, the finding stress the importance of breaking the stereotypical dogma of Mendelian inheritance in “monogenic” diseases and emphasizes the importance of large scale genome-wide case-control genetic studies in elucidating the genetic causes of inherited diseases largely unsolved by sequencing approaches.

In conclusion, this study provides a novel GWAS-based framework for systematically detecting disease-associated variants, unbiased with regard to genomic location and mode of genetic influence, in so-called “monogenic” disorders. It also highlights the under-appreciated significance of non-Mendelian high frequency variants that may significantly account for the undetermined heritability of various inherited diseases. At the same time, significance of these variants may extend beyond genetic diagnosis as they may simultaneously serve as ideal targets of local genome treatments.

## Methods

### Patients and controls

The study was initiated after ethical approvals were granted by the Institutional Review Boards of Kyushu University Hospital, Tohoku University Hospital Yuko Wada Eye Clinic, Nagoya University Hospital, and Juntendo University Hospital, from which 944 presumed-unrelated patients with RP were recruited. The study also received approval from Tokyo Medical and Dental University. The majority of the patients were recruited through a genetic screening project hosted by the Japan Retinitis Pigmentosa Registry Project (JRPRP) in which 83 genes associated with RP were analyzed by targeted re-sequencing^11^. The remaining patients were recruited from Tohoku University Hospital. Most of the unaffected controls, who were ruled out for RP with a fundus examination, were recruited at Tohoku University Hospital and its affiliated hospitals^39^. The remaining control samples from subjects with no documented history of ocular disease were purchased from the National Institutes of Biomedical Innovation, Health and Nutrition (https://bioresource.nibiohn.go.jp/). All procedures followed the tenets of the Declaration of Helsinki. Informed consent was obtained from all patients and controls before collecting blood samples for DNA extraction and establishing patient-derived lymphoblastoid cell lines (LCLs).

### Genome-wide association study

In the first GWAS, 644 cases and 620 controls, all from Japan, were genotyped with the CoreExome-24 v1.1 (Illumina, San Diego, CA, USA). The total number of analyzed samples was reduced to 581 cases and 603 controls after quality control (QC). During QC, we excluded single nucleotide variants (SNVs) with Hardy-Weinberg equilibrium (HWE) *P* < 0.0001 in the controls, a call rate < 99%, or three alleles. Data were also discarded if the sample had a call rate < 98%. In addition, closely related pairs (pi-hat >0.1)^40^, or ancestral outliers, as determined with a PCA analysis using the 1000 Genomes Project (five Asians, CEU, and YRI) and PLINK software were removed. One hundred and forty-nine cases with causal mutations identified after targeted re-sequencing^11^ were also removed. The remaining 432 cases and 603 controls were subjected to a GWAS using 10,673,864 variants following whole-genome imputation of 523,187 genotyped SNVs using phased haplotypes from the 1000 Genomes Project (Phase 3) as the reference panel. SHAPEIT was used for phasing, followed by minimac3 for genotype imputations^41^. Imputed variants with estimated imputation accuracy of Rsq > 0.3 were selected. It should be noted that the variants were not excluded based on minor allele frequency (MAF) in this study because assumed rare RP mutations may be tagged better with lower frequency variants. Statistical analysis of the GWAS was performed using RVtests^42^. We used imputed genotype dosages and top 10 principle components as covariates for the analysis input data. The principal component scores were calculated using PLINK. The Wald test was used as the association model in the analysis.

The second GWAS comprised 300 cases and 300 controls and was also carried out using genotyping with the CoreExome-24 v1.2 (Illumina). The total number of samples was reduced to 286 cases and 287 controls after applying QC procedures identical to the first GWAS. Samples were also removed if they overlapped with the first GWAS. Then, 78 cases with causal mutations identified through targeted re-sequencing were excluded^11^. The remaining 208 cases and 287 controls were subjected to a GWAS using 10,383,808 SNVs following whole-genome imputation of 522,207 genotyped SNVs selected with the same criteria as the first GWAS.

A meta-analysis combining the first and second GWAS data sets was performed using METAL^43^. Stepwise conditional analyses, starting with a top associated variant, were performed with the dosages of target variants of the regions used as covariates. Any variant at *P* < 10^−5^ was then assumed to be independent from the main signal in the region.

To assess linkage between nonsynonymous variants identified through a previous targeted re-sequencing study^11^ and the GWAS peak variants positioned within 1Mb of each other, correlation coefficients (r^2^) and D’ / LOD were calculated using Haploview.

### WGS and Sanger sequencing

We performed WGS using the NovaSeq 6000 (Illumina, San Diego, CA, USA) sequencer with 151 bp paired-end reads. The sequencing library was constructed using the TruSeq Nano DNA Library Prep Kit (Illumina) according to the manufacturer’s instructions. The sequenced reads were aligned to the human reference genome using BWA-mem (ver. 0.7.17). Then, PCR duplicate reads were marked using Picard tools (ver. 2.17.8). Base quality scores were recalibrated, and SNVs and short insertions and deletions were called, using GATK (ver. 4.1.2.0) according to the GATK Best Practices (https://software.broadinstitute.org/gatk/best-practices/). Structural variants were called using Manta^44^ according to the instructions and with default parameters. In addition, we used IGV software to visually inspect reads for specific genes that were reported to carry structural variants.

Sanger sequencing was carried out for genotyping of family members using the protocol described earlier^45^. In brief, genomic DNA was amplified with PCR using Amplitaq Gold and a primer pair designed by Primer3 (ver. 0.4.0; http://bioinfo.ut.ee/primer3-0.4.0/). PCR amplification was performed in a 20 μl total volume containing 20 ng genomic DNA, 1x GoTaq buffer, 0.5 mM dNTPs, 10 μM of each primer, and 2 units (5 U/μl) of GoTaq polymerase (Promega, Madison, Wisconsin). The PCR amplicons were applied onto a 2% agarose gel with appropriate controls and markers.

### mRNA analysis using patient-derived lymphoblastoid cell lines (LCLs)

To generate patient-derived LCLs, lymphocytes were transformed with the Epstein-Barr virus at a core facility run by Tokyo Medical Dental University. gRNAs were designed and T7E1 assay were performed as previously described (Figure S3)^15^. A plasmid for CAG promoter insertion genome editing was constructed as described previously (Figure S3)^15^. The donor sequence included a CMV promoter (from pCAG-Neo, Wako, Osaka, Japan) for in-frame insertion upstream of the *EYS* start codon. The donor template, which comprised the flanking micro-homology arms, gRNA target sites (5’GTCCAATTTACCACATATGATGAGGGT3’) and the donor sequence, was sub-cloned and inserted into the vector (pX601, addgene #61591) using a DNA ligation kit (Clontech, Mountain View, CA). A plasmid for mutation replacement genome editing was constructed as described previously^15^. The donner template, which comprised the flanking micro-homology arms, gRNA-1 target site or gRNA-4 target site and the donor sequence, were sub-cloned and inserted into the vector (pX601, addgene #61591) using a DNA ligation kit (Clontech, Mountain View, CA). To avoid repeated cleavage after mutation replacement, mutations were introduced in the flanking gRNA target sites. The mutation in the 5’ gRNA-1 and 3’ gRNA-4 target sites were selected using codon optimization tool GENEisu (http://www.geneius.de/GENEius/) on human codon table. The LCLs were transfected with a plasmid using Trans-IT XP transfection reagent (Mirus Bio, Madison, WI) and treated with a demethylating agent, 5-Aza-2’-deoxycytidine (1 μM; Abcam, Cambridge, UK), and hydralazine hydrochloride (0.2 μM; Abcam). To test if transcripts were degraded by nonsense-mediated mRNA decay(NMD), LCL were treated by emetine (Sigma-Aldrich, St. Louis, MO) at 60 μg/mL for 12hrs before RNA extraction^46^. For mutation replacement gene editing, LCL was co-transfected with the CAG promoter insertion plasmid and the mutation replacement genome editing plasmid (ratio 1:3). Total RNA was extracted 48 hrs post-transfection using the miRNeasy plus mini kit (Qiagen, Hilden, Germany) according to the manufacturer’s instructions. A 500-ng sample of total RNA was reverse-transcribed with SuperScript IV (Thermo Fisher Scientific, Waltham, MA) and oligo(dT) primers (Thermo Fisher Scientific) at 55° C for 30 min. The design of the primer sets for RT-PCR is shown in Table S6. The RT-PCR reaction was performed with KOD One DNA polymerase (Toyobo, Osaka, Japan). PCR products were analyzed on agarose gels.

#### Zebrafish experiments

Zebrafish (Danio rerio) AB strain was maintained and bred in a standard fashion^47^. Morpholino oligonucleotides and eys mRNA microinjection were performed as described previously^48, 49^. Morpholino oligonucleotides were obtained from Gene Tools LLC. The following morpholinos were used: ATG-MO 5’-CTCATGTTTGTCTTGGCTCGACTGG-3’; SP-MO, 5’-TTGACTTACCCTTAAATCCTGGTG-3’; Standard Control Morpholino, 5’-CCTCTTACCTCAGTTACAATTTATA-3’. Human EYS cDNAs, wildtype and c.2528G>A were cloned into pcDNA 3.1(+) vector and were transcribed by the mMessage Machine T7 kit (Ambion®, Thermo Fisher Scientific, Waltham, Massachusetts, U.S.A.). In morpholino knockdown experiments with or without rescue mRNA, mixture of 200 mM ATG-MO, SP-MO and 300ng/ul mRNA or 380 mM ATG-MO was applied, respectively. We injected morphorinos and mRNA into embryos within 40 minutes after fertilization

After sacrificing the fish, zebrafish embryos or adult head were placed in 4% (w/v) paraformaldehyde (PFA), pH 7.4, in PBS overnight at 4 °C, and then incubated in 20% glucose aqueous solution overnight at 4 °C. The fixed fish were embedded in optimal cutting temperature compound (Neg50TM, Thermo Fisher Scientific) and quick-frozen in −150 °C nitrogen freezer. The samples were cut into 14-16 µm sections. The sections were placed in a solution containing 0.1 M PBS, 10 % BSA for 2 hr at room temperature, and then incubated with the primary antibody at 4 °C overnight. After a 0.1 M PBS/0.005% Tween rinse cycle, the sections were incubated with Alexa Fluor 594-conjugated IgG antibodies (1:500, Jackson ImmunoResearch, West Grove, Pennsylvania, U.S.A.) species-matched for the primary antibody, rhodamine-conjugated phalloidin (1:200, Cytoskeleton Denver, Colorado, U.S.A.), or DAPI (1:1,000, Cytoskeleton) at room temperature for 2 h. The slides were then mounted with an aqueous mounting medium (PermaFluor®, Thermo Fisher Scientific) after another cycle of rinsing. The sections were analyzed with the use of the confocal fluorescence laser microscopes LSM 710 (Carl Zeiss, Jena, Thüringen, Germany). The following primary antibodies and dilutions were used: rabbit polyclonal anti-Eys antibody (1:200, Novus Biologicals, Centennial, Colorado, USA), rabbit polyclonal anti-rhodopsin (bovine) antibody (1:500, Abcam, Cambridge United Kingdom), and mouse monoclonal [6-11B-1] anti-alpha Tubulin (acetyl K40) antibody (1:200, Abcam, Cambridge United Kingdom).

### Statistics

The frequency of homozygotes of the G843E and S1653fs mutations was calculated as the square of allele frequency of each mutation in the general population, assuming random mating. Fisher’s exact test was carried out using JMP software (SAS Institute, USA) to assess the significance of the enrichment of the G843E mutation.

## Acknowledgements

The study was supported by the Japan Retinitis Pigmentosa Registry Project. We thank J. Inazawa and M. Takaoka (Bioresource Laboratory, Tokyo Medical and Dental University) for technical support in establishing the patient-derived LCLs.

## Authors’ contributions

KMN designed the overall study. Collection of DNA samples and clinical assessment was performed by KMN, HK, TA, YW, SU, and YI, and was led by AM, HT, KHS, TI and TN. FM, YK, KC, JN, MA and TK performed the bioinformatics analyses. KF and MT performed the LCL studies. Zebrafish experiments were carried out by YM, TT, and MT. Sequencing was carried out by FM, KF, MT, and YM. KMN, CR, TT, and TN provided the funding.

All authors read and approved the final manuscript.

## Funding

This study was supported in part by the Japan Agency for Medical Research and Development (#19ek0109213h0003, KMN and #JP18lk1403004, TA).

## Availability of data and materials

The datasets generated during and/or analyzed during the current study are available from the corresponding author on reasonable request.

## Ethics approval and consent to participate

The study was initiated after ethical approvals were granted by the Institutional Review Boards of Kyushu University Hospital, Tohoku University Hospital, the Yuko Wada Eye Clinic, Tokyo Medical and Dental University, and Nagoya University Hospital. All procedures followed the tenets of the Declaration of Helsinki. Informed consent was obtained from all patients and controls before collecting blood samples for DNA extraction and establishing patient-derived LCLs.

## Competing interests

The authors declare that they have no competing interests.

## Supplementary Figures

**Figure S1.**
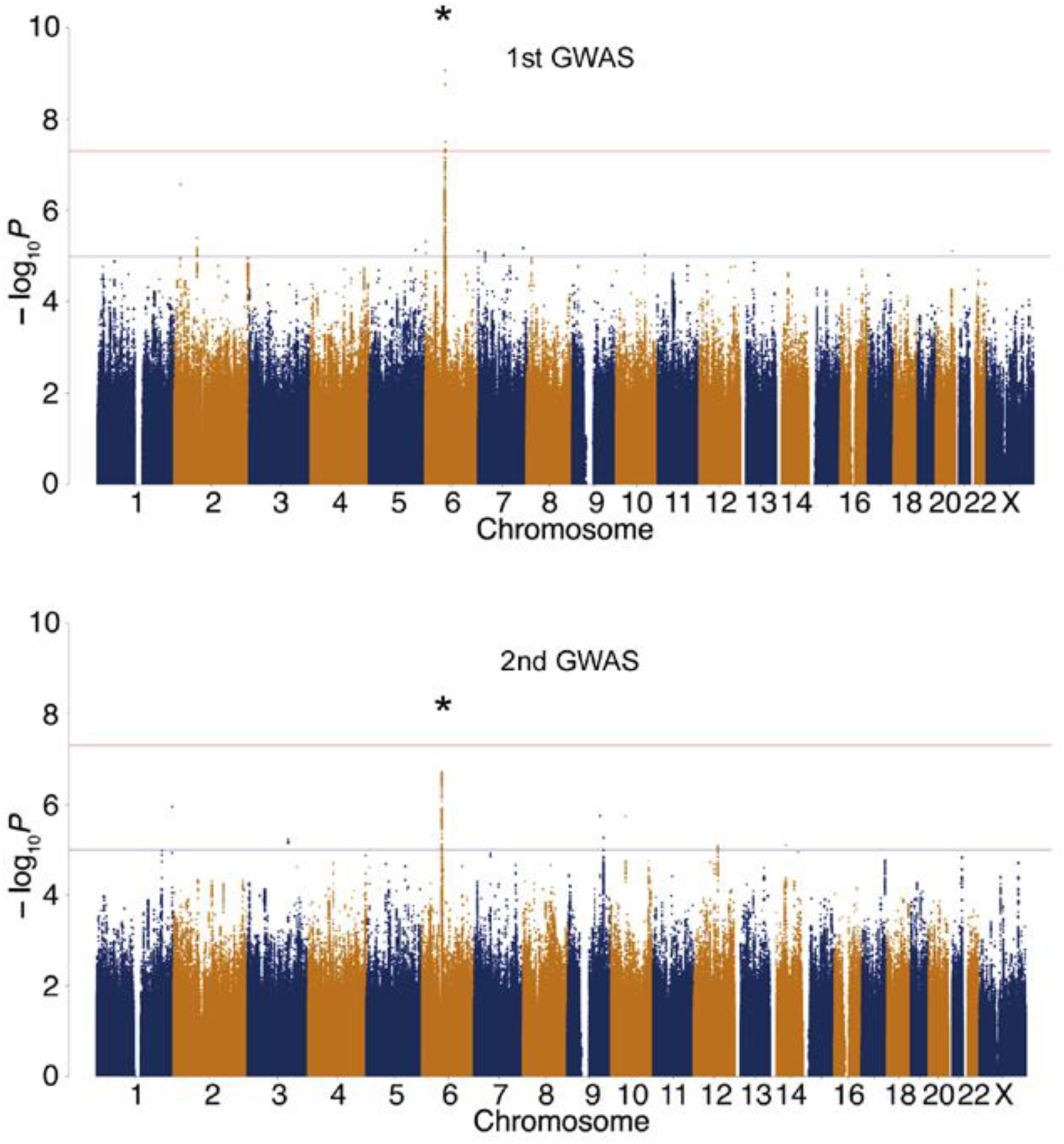
Manhattan plots of 1^st^ GWAS and 2^nd^ GWAS. The *EYS* locus is labeled with an asterisk. GWAS, genome-wide association study. Genome-wide significance (*P* = 5.0 ×10^-8^) and possible significance (*P* = 1.0×10^-5^) are marked with red and blue lines, respectively.

**Figure S2.**
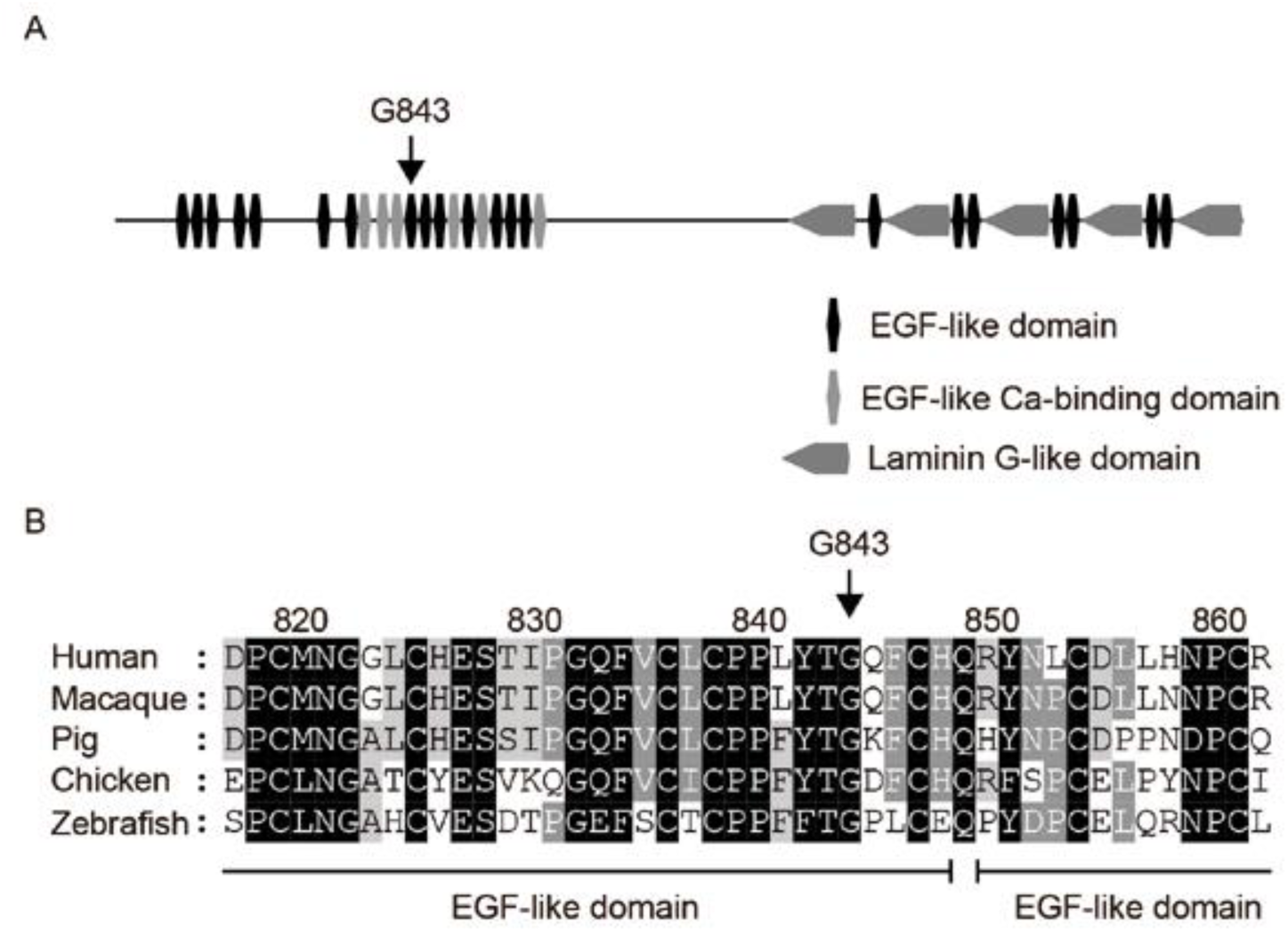
Conservation of G843 across diverse species and *in silico* analysis. **A**. Domain structure of *EYS* in relation to G843E. **B**. Conservation of G843 across diverse species ranging from zebrafish to humans. The multiple sequence alignment was generated using ClustalW (https://clustalw.ddbj.nig.ac.jp/). Accession numbers of the protein sequences used for sequence comparison are as follows: human, NM_001142800.1; macaque, XM_011737495.1; pig, XM_021084496; chicken, XM_015284845.1; zebrafish, XM_009307513.

**Figure S3.**
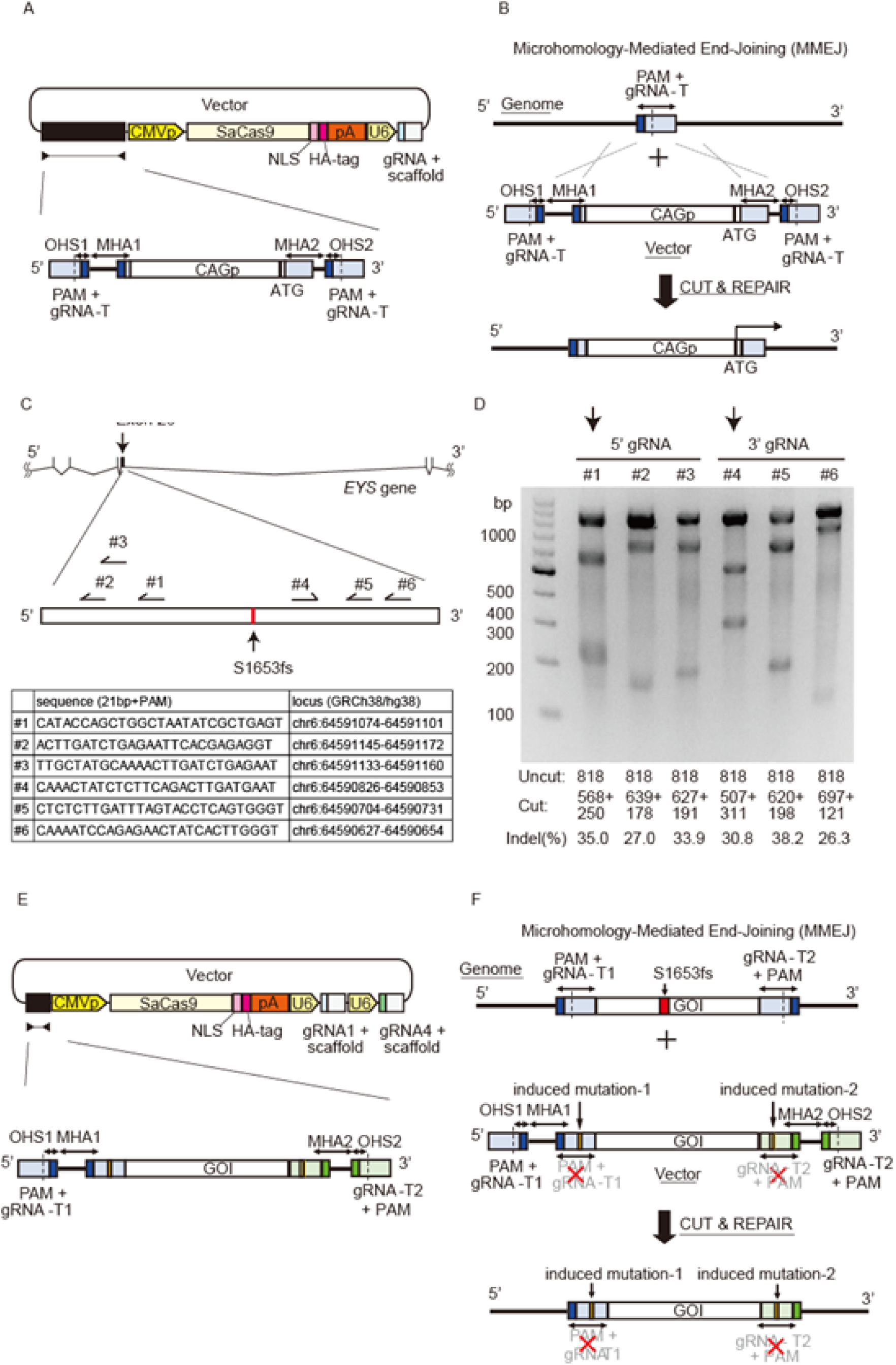
Design of plasmid and strategy to edit patient-derived lymphoblastoid cell lines (LCLs) **A**. Design of targeting plasmid to insert a *CAG* promoter upstream of the *EYS* gene through genome editing. **B**. Illustration of the microhomology-mediated end-joining *CAG* promoter knock-in strategy. The DNA sequence of the LCL genome and plasmid vector were excised at the gRNA target sites (gRNA-T; dotted line) by two gRNAs and SaCas9. A *CAG* promoter was inserted into the genome using micro homology arms (MHA). **C**. Schematic map of gRNA designed inside exon 26 and list of their sequences. **D**. T7E1 assay for each gRNAs. Expected DNA size and quantified editing efficiency are displayed under the representative gel image. gRNA-1 and -4 were selected for the downstream mutation replacement genome editing experiment. **E**. Design of targeting plasmid to mutation replacement genome editing of *EYS* gene through genome editing. **F**. Illustration of microhomology-mediated end-joining mutation replacement strategy. Genome of interest (GOI) with and without the S1653Kfs mutation are excised at the flanking gRNA target sites (gRNA-T; dotted line) from LCL genome and plasmid vector, respectively, by two gRNAs (1 and 4) and SaCas9. GOI without S1653Kfs mutation is inserted into the genome using MHA. gRNA-T, guide RNA target; PAM, protospacer adjacent motif; OHS, over-hanging sequence; NLS, nuclear localizing signal; pA, ploy A; U6, human U6 promoter.

**Figure S4.**
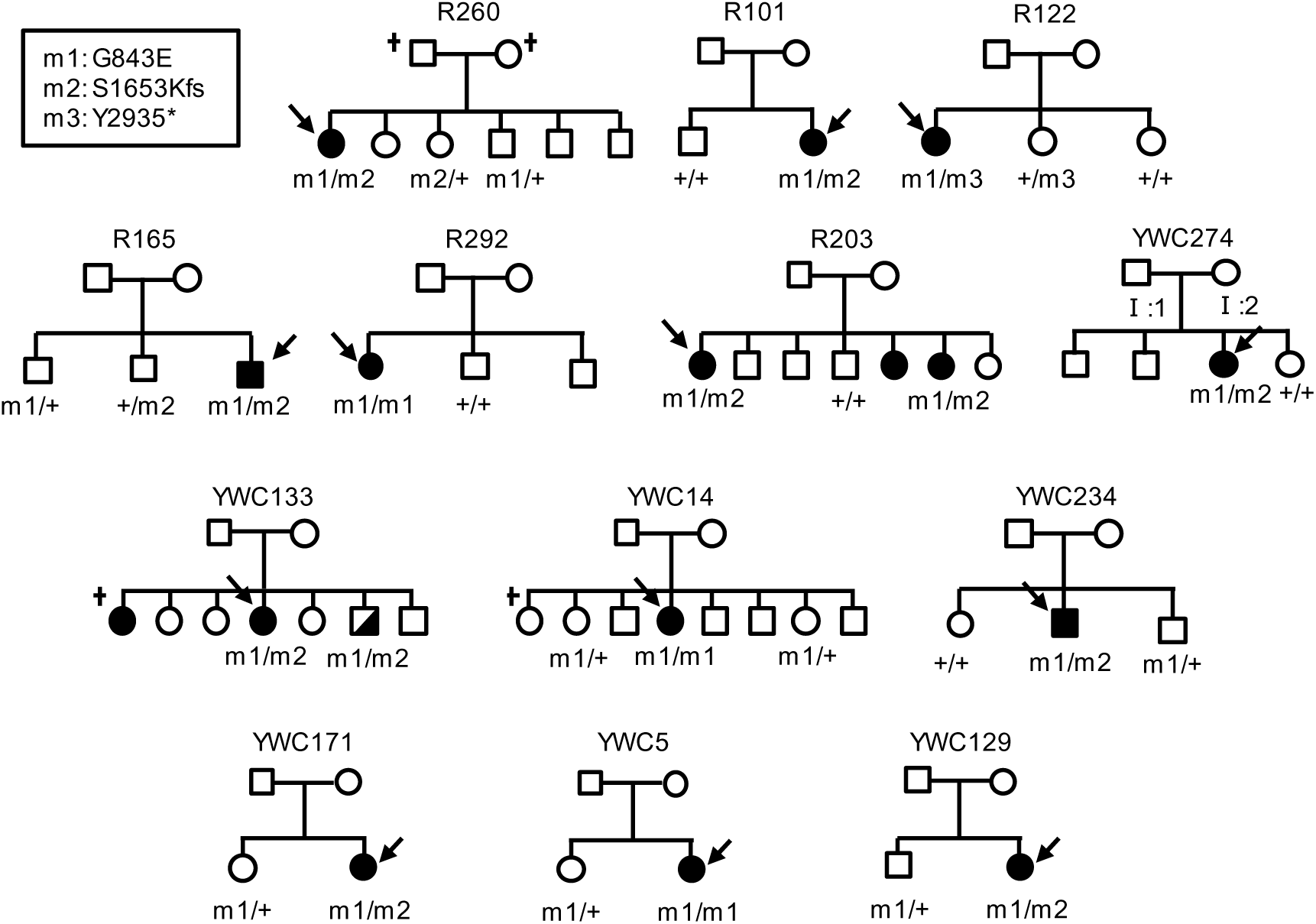
Co-segregation of the G843E genotype with phenotype as an ARRP allele. The arrows indicate index patients. The half-filled symbol indicates an asymptomatic RP patient initially diagnosed as unaffected.

**Table S1.** Summary of 1^st^ and 2^nd^ GWAS. GWAS, genome-wide association study; No., number; QC, quality control; SNV, single nucleotide variant.

|  | 1st GWAS | 2nd GWAS |
| --- | --- | --- |
| No. of <u>cases/controls</u> genotyped | 644/624 | 300/300 |
| No. of <u>cases/controls</u> after QC | 581/603 | 286/287 |
| No. of <u>cases</u> solved by sequencing | 149 | 78 |
| No. of <u>cases/controls</u> used in GWAS | 432/603 | 208/287 |
| No. of genotyped SNV used | 523,187 | 522,207 |
| No. of SNV after imputation | 10,673,864 | 10,383,808 |
| $\lambda$ | 1.061 | 1.037 |

**Table S2.** List of candidate loci at *P* < 1.0 ×10^-5^ after 1st GWAS. Chr, chromosome; Ref, reference sequence; Alt, alternative sequence; AF, allele frequency.

| Gene_name | Genomic location | Chr | Coordinate | Ref | Alt | ToMMo AF | Odds ratio | P-value |
| --- | --- | --- | --- | --- | --- | --- | --- | --- |
| NT5C1B-RDH14(dist=58909),LINC013<br>76(dist=337974) | intergenic | 2 | 18829755 | A | T | 0.073 | 0.128 | 2.70E-07 |
| EGR4(dist=3282),ALMS1(dis<br>t=88775) | intergenic | 2 | 73524111 | G | GGT | ND | 1.723 | 3.93E-06 |
| CAMK2A | intronic | 5 | 149610967 | AAAT | A | ND | 0.558 | 7.37E-06 |
| EYS | intronic | 6 | 65728469 | C | T | ND | 4.005 | 8.61E-10 |
| DNAAF5 | intronic | 7 | 791207 | ATT | A | ND | 1.621 | 7.71E-06 |
| STK31(dist=421431),NPY(di<br>st=30246) | intergenic | 7 | 24293561 | G | A | 0.140 | 1.914 | 8.27E-06 |
| SEMA3A(dist=231370),LOC1<br>01927378(dist=106205) | intergenic | 7 | 84055587 | T | TATA<br>G | ND | 0.494 | 9.55E-06 |
| NOC3L | exonic<br>(:c.A1415C:p.E472A) | 10 | 96104665 | T | G | 0.305 | 0.632 | 9.12E-06 |

**Table S3.**
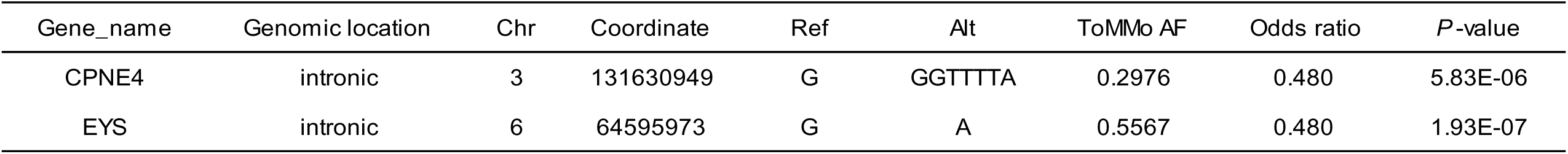
List of candidate loci at *P* < 1.0 ×10^-5^ after 2nd GWAS. Chr, chromosome; Ref, reference sequence; Alt, alternative sequence; AF, allele frequency.

**Table S4.**
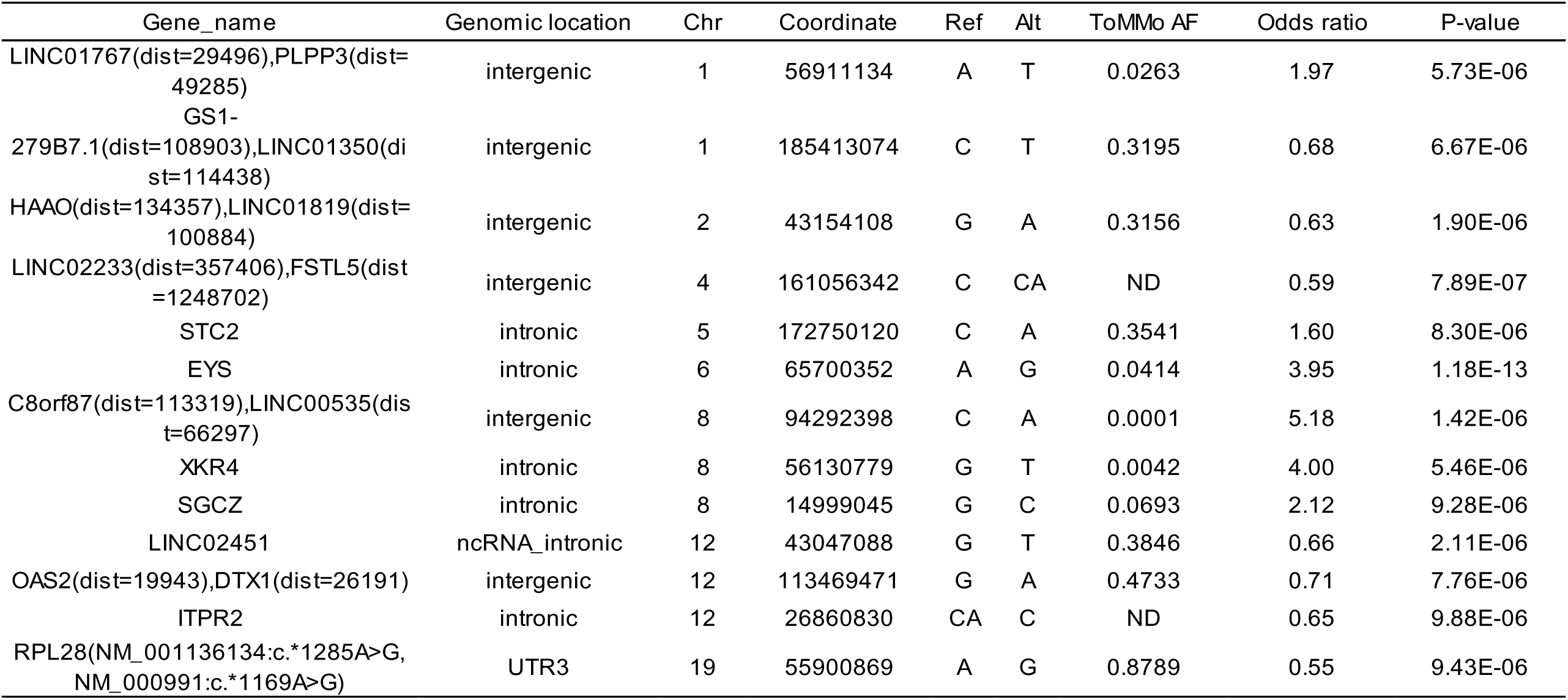
List of candidate loci at *P* < 1.0 ×10^-5^ after meta GWAS. Chr, chromosome; Ref, reference sequence; Alt, alternative sequence; AF, allele frequency.

**Table S5.**
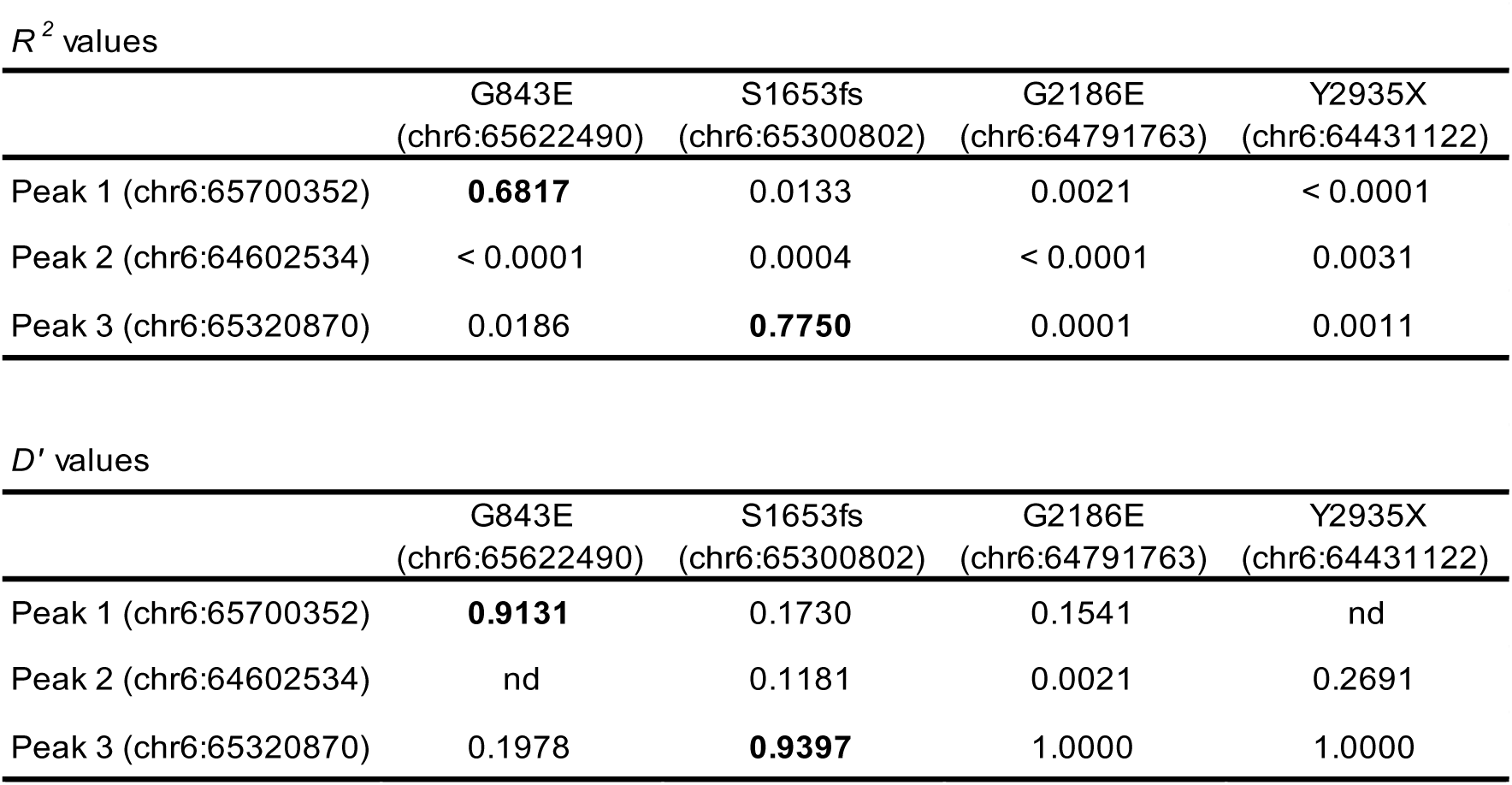
Evaluation of LD of top SNPs for *EYS* peaks in RP patients.

|  | G843E<br>(chr6:65622490) | S1653fs<br>(chr6:65300802) | G2186E<br>(chr6:64791763) | Y2935X<br>(chr6:64431122) |
| --- | --- | --- | --- | --- |
| Peak 1 (chr6:65700352) | **0.6817** | 0.0133 | 0.0021 | < 0.0001 |
| Peak 2 (chr6:64602534) | < 0.0001 | 0.0004 | < 0.0001 | 0.0031 |
| Peak 3 (chr6:65320870) | 0.0186 | **0.7750** | 0.0001 | 0.0011 |

**Table S5 Evaluation of LD of top SNPs for *EYS* peaks in RP patients**
|  | G843E<br>(chr6:65622490) | S1653fs<br>(chr6:65300802) | G2186E<br>(chr6:64791763) | Y2935X<br>(chr6:64431122) |
| --- | --- | --- | --- | --- |
| Peak 1 (chr6:65700352) | **0.9131** | 0.1730 | 0.1541 | nd |
| Peak 2 (chr6:64602534) | nd | 0.1181 | 0.0021 | 0.2691 |
| Peak 3 (chr6:65320870) | 0.1978 | **0.9397** | 1.0000 | 1.0000 |

**Table S6.**
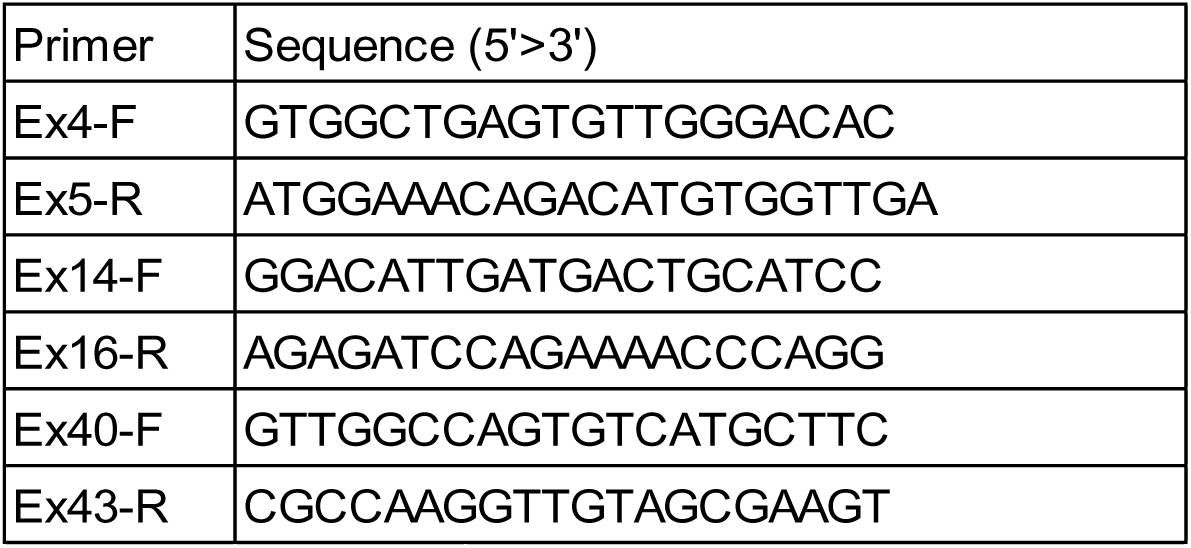
Primers used for RT-PCR.

## Notes

This study has been supported in part by the Japan Agency for Medical Research and Development (#18ek0109213h0001, KMN and #JP18lk1403004, TA).

